# Screening of candidate substrates and coupling ions of transporters by thermostability shift assays

**DOI:** 10.1101/367805

**Authors:** Homa Majd, Martin S. King, Shane M. Palmer, Anthony C. Smith, Liam D.H. Elbourne, Ian T. Paulsen, David Sharples, Peter J. F. Henderson, Edmund R. S. Kunji

## Abstract

Substrates of most transport proteins have not been identified, limiting our understanding of their role in physiology and disease. Traditional identification methods use transport assays with radioactive compounds, but they are technically challenging and many compounds are unavailable in radioactive form or are prohibitively expensive, precluding large-scale trials. Here, we present a high-throughput screening method that can identify candidate substrates from libraries of unlabeled compounds. The assay is based on the principle that transport proteins recognize substrates through specific interactions, which lead to enhanced stabilization of the transporter population in thermostability shift assays. Representatives of three different transporter (super)families were tested, which differ in structure as well as transport and ion coupling mechanisms. In each case, the substrates were identified correctly from a large set of chemically related compounds, including stereo-isoforms. In some cases, stabilization by substrate binding was enhanced further by ions, providing testable hypotheses on energy coupling mechanisms.

## Introduction

Transport proteins, also called transporters, porters, carriers, translocases or permeases, encompass a diverse and ubiquitous group of membrane proteins that facilitate the translocation of ions and molecules across all types of biological membranes, linking biochemical pathways, maintaining homeostasis and providing building blocks for growth and maintenance. They comprise 5-15% of the genomic complement in most organisms. Transport proteins are classified into four major groups; primary active transporters, secondary active transporters, facilitative transporters, and channels. Human solute transport proteins have been divided into 65 subfamilies (based on sequence alignments or experimentally determined substrate specificities), which transport a diverse range of compounds, including amino acids, sugars, nucleotides, lipids, vitamins, hormones, inorganic and organic ions, metals, xenobiotics and drugs (http://slc.bioparadigms.org/). Of the 538 identified solute transport proteins in humans, more than a quarter have no assigned substrate specificity (http://slc.bioparadigms.org/). Consequently, their role in human physiology is ill-defined and opportunities for drug intervention are missed (Hediger et al., 2013). Currently, about half of the transport proteins have been associated with human disease (Rask-Andersen et al., 2013; Hediger et al., 2013; Lin et al., 2015), but only 12 classes of drugs target them directly, despite their inherent ‘druggability’ (Hediger et al., 2013; Lin et al., 2015; Cesar-Razquin et al., 2015; Fauman et al., 2011). The situation is not better in other eukaryotic organisms and bacterial pathogens, currently on the WHO list (**Supplementary Table 1**) (Elbourne et al., 2017). For most transporters in the sequence databases the identifications are preliminary because they are based only on sequence homology without direct experimental evidence for the substrates, even though single amino acid residue variations in the substrate binding site can alter the specificity profoundly. Moreover, the substrate specificities of some transporters may have been incompletely or incorrectly assigned. Finally, there could be membrane proteins with unassigned function that belong to unidentified transporter families, which are not counted at all.

One of the major challenges is to find the correct substrates from a large number of potential candidates. There are approximately ˜3,000 metabolites in *E. coli* (Sajed et al., 2016), ˜16,000 in *S. cerevisiae* (Ramirez-Gaona et al., 2017), ˜40,000 in *Homo sapiens* (Wishart et al., 2013), and plants must have even more as they carry out an extensive secondary metabolism. Some metabolites, such as vitamin B_6_, have several interconvertible species, each of which could be transported. The classical method to identify transport proteins monitors the accumulation of radiolabeled compounds in whole cells, membrane vesicles or proteoliposomes. However, these experiments can easily fail when the expressed transporters are not active due to targeting, insertion or folding issues, when they are unstable in purification and reconstitution experiments, or when substrate and coupling ion gradients are not setup correctly. Moreover, some compounds are not available in radioactive form or are prohibitively expensive, preventing large-scale identification trials. Given all these technical difficulties, it is often necessary to limit the number of candidate substrates first by using phylogenetic analysis, by analyzing phenotypic (patho)physiological data, by complementation studies or by metabolic analysis of knock-out or mutant strains.

Therefore, there is an unmet demand for the development of new methods to limit the number of potential substrates for identification of solute carriers. Here, we present a high-throughput screening method for the identification of substrates of transporters, which does not require radioactive compounds or prior knowledge. The method uses the simple concept that transporters recognize their substrates through specific interactions, enhancing their stabilization in thermostability shift assays. We verify the approach by defining the substrate specificity of three solute carriers from different bacterial and eukaryotic protein families and show that these experiments also provide valuable clues about the ion coupling mechanism.

## Results

### Principle of the method

Ligands, such as substrates or inhibitors, are recognized by transport proteins through specific interactions at the exclusion of other molecules. The formation of these additional bonds leads to an increase in the total number of interactions (Robinson and Kunji, 2006; Yan, 2017; Yamashita et al., 2005). Consequently, binding leads to an overall increase in the stability of the ligand-bound species compared to the unliganded species in the population of protein molecules. We have previously shown that the mitochondrial ADP/ATP carrier (AAC) from the thermophilic fungus *Thermothelomyces thermophila* (UniProt G2QNH0) when purified in dodecyl-maltoside is stabilized upon binding of its specific inhibitors carboxyatractyloside and bongkrekic acid in thermostability shift assays using the thiol-reactive fluorophore N-[4- (7-diethylamino-4-methyl-3-coumarinyl)-phenyl]-maleimide (CPM) (Crichton et al., 2015; King et al., 2016). In the assay, the apparent melting temperature T_m_ of a population of purified transporters is determined by monitoring the increase in fluorescence by CPM reacting with thiols that have become exposed due to thermal denaturation of the proteins (**Fig. 1a**). The apparent melting temperature T_m_ is the temperature at which the rate of unfolding for a given population is highest. We tested whether transported substrates can also cause a shift in thermostability, as their properties differ quite substantially from those of inhibitors or other tight binders. Transported substrates bind only transiently and relatively weakly, leading to a conformational change that in turn causes the release of the substrate on the other side of the membrane, so it was not obvious that this approach would work. However, the thermostability of the AAC population was enhanced in the presence of ADP and ATP, but not in the presence of AMP, which reflects the known substrate specificity of the carrier well (Mifsud et al., 2013) (**Fig. 1b**). This effect is only observed at high concentrations of the substrate, well above the apparent Km of transport (**Supplementary Fig. 1**). The higher the substrate concentration, the higher the likelihood that part of the population is prevented from unfolding by binding of the substrate, leading to the observed shift in thermostability. We reasoned that this approach could be applied as a screening method to find substrate candidates of uncharacterized transporters by using compound libraries. To verify the method, we have tested transporters from three different (super)families, which are distinct in structure and transport mechanism.

**Figure 1.**
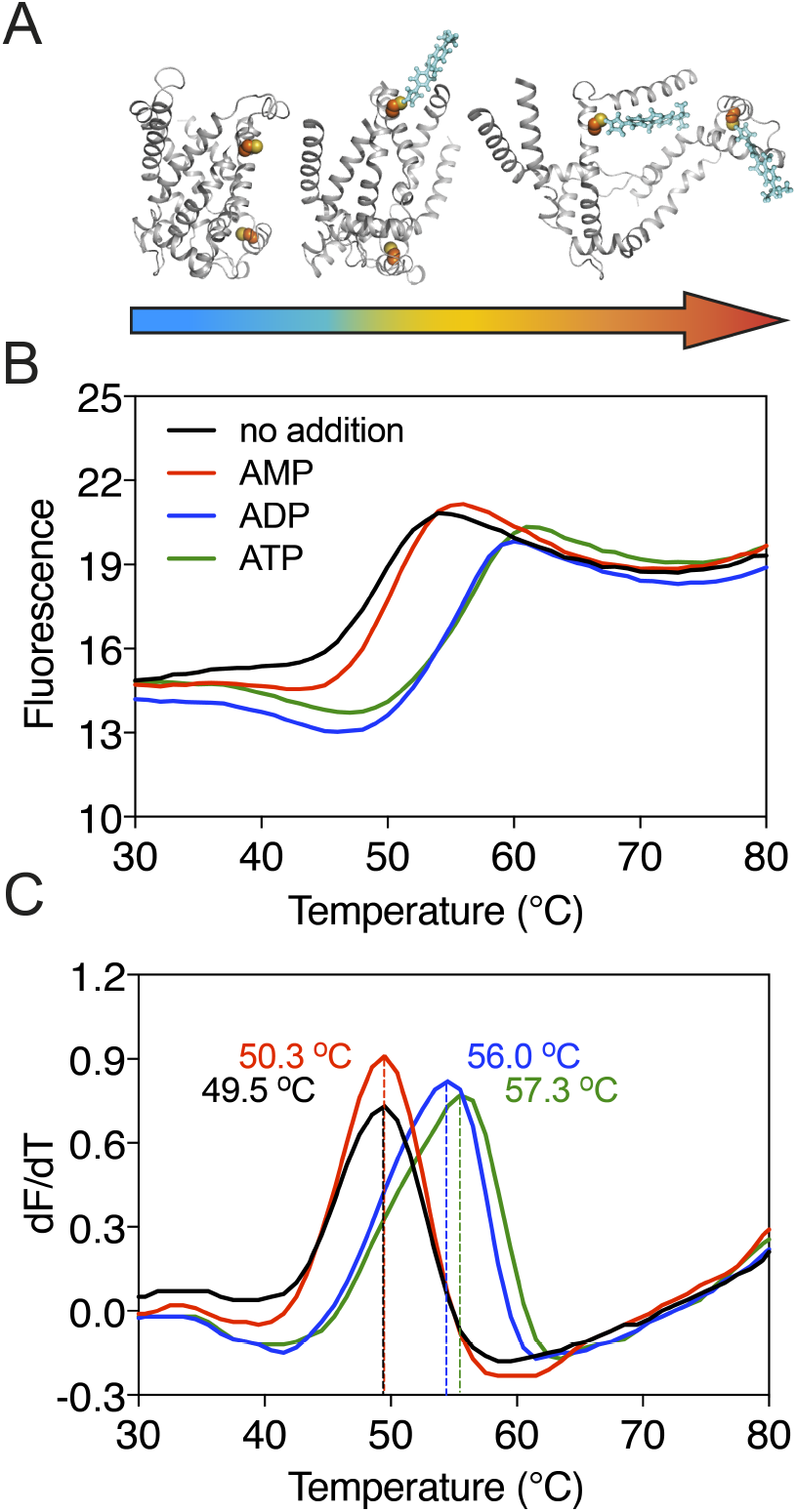
Substrate-induced stabilization of a mitochondrial ADP/ATP carrier. (**a**) As protein molecules in a population unfold due to a gradual rise in temperature (25–90 °C), buried cysteine residues become solvent-exposed and accessible to the thiol-specific probe CPM (blue stick representation) that becomes fluorescent upon reaction. (**b**) Typical unfolding curves of the mitochondrial ADP/ATP carrier of *Thermothelomyces thermophila (*2 µg) in the absence and presence of AMP, ADP and ATP, shown in red, blue and green, respectively. (**c**) The apparent melting temperature (T_m_) is the peak in the derivative of the unfolding curve (dF/dT) (below), which is used as an indicator of thermal stability. The apparent melting temperatures reported in the text are from three independent protein purifications.

### Substrates cause specific thermostability shifts in different transporters

The galactose transporter GalP of *E. coli* (Henderson, 1977) is a prototypical member of the sugar porter family that belongs to the major facilitator superfamily (MFS) (Pao et al., 1998). Currently, the MFS contains 24 different transporter families and 320,665 sequence entries from 5224 species, but the substrates for the vast majority of them have not been formally identified (Pfam CL0015). In humans, 14 of the 65 solute carrier families belong to the MFS, and substrates are not known for around 12% of them (http://slc.bioparadigms.org/).

The structure of GalP has not been determined, but those of its mammalian homologs GLUT1 (Deng et al., 2014), GLUT3 (Deng et al., 2015), and GLUT5 (Nomura et al., 2015) are available (**Fig. 2a**). GalP contains three cysteine residues, of which only one is readily accessible to reaction with *N*-ethylmaleimide (McDonald and Henderson, 2001). To evaluate the strategy, we measured the unfolding curves of purified GalP in dodecyl-maltoside in the presence of a large number of different sugars. We determined the temperature shift by subtracting the apparent T_m_ of unliganded GalP (57.6 ± 0.3 °C) from the apparent T_m_ values observed in the presence of compounds (ΔT_m_) (**Supplementary Fig. 2**). The GalP population was markedly stabilized by D-glucosamine (ΔT_m_; 5.7 ± 0.4 °C), D-glucose (ΔT_m_; 4.2 ± 0.5 °C), D-galactose (ΔT_m_; 2.1 ± 0.6 °C), and to a lesser extent by D-talose, 2-deoxy-D-glucose, 6-deoxy-D-glucose. Importantly, the related L-isoforms showed no significant shift (**Fig. 2b**). These results match the known substrate specificity of GalP well (Henderson and Maiden, 1990; Walmsley et al., 1994). D-Glucosamine, the most stabilizing compound, had not been investigated previously, but transport assays showed that this compound is a new substrate (**Supplementary Fig. 3**), demonstrating that the assay can be used to discover as well as to extend the substrate specificity of transport proteins.

**Figure 2.**
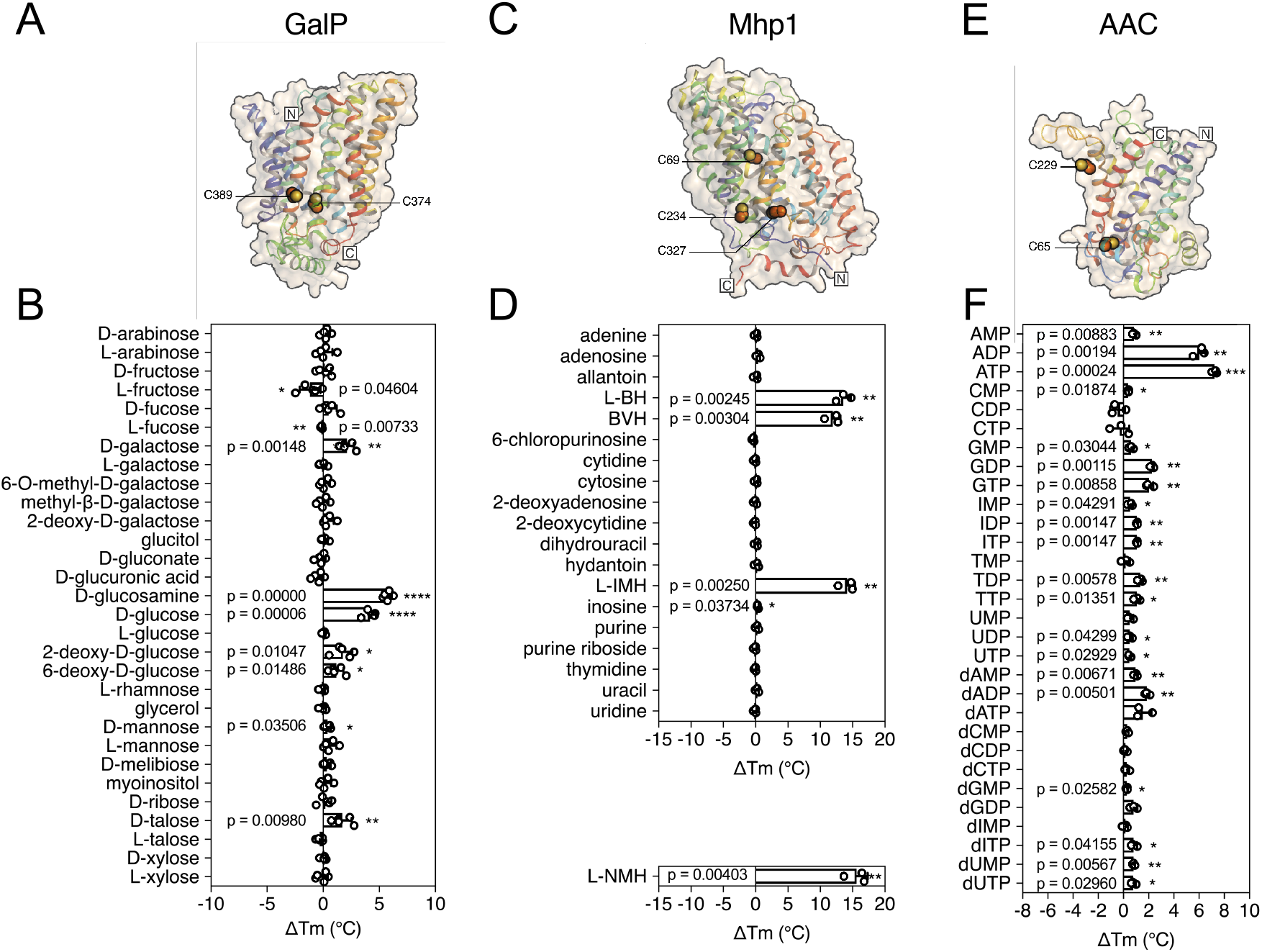
Validation of the method for determining substrate specificity using three unrelated proteins. (**a**), (**c**) and (**e**) Structural models of GalP (based on GLUT5, PDB 4YBQ), Mhp1 (PDB 2JLN) and AAC (based on Aac2p, PDB 4C9G), respectively. The models are shown in rainbow cartoon and wheat surface representations. Cysteine residues are shown as spheres, except for Cys-19 of GalP, which could not be modeled. (**b**), (**d**) and (**f**) Thermostability screen of GalP (3 µg), Mhp1 (3 µg) and AAC (2 µg) against sugar, nucleobase and nucleotide libraries, respectively. The temperature shift (ΔT_m_) is the apparent melting temperature in the presence of compound minus the apparent melting temperature in the absence of compound. The data are represented by the standard deviation of five, three and three independent repeats, respectively. Two-tailed Student’s *t*-tests assuming unequal variances were performed for the significance analysis (0.05 <*p*-value: not significant; 0.01<*p*-value<0.05: *; 0.001<*p*-value<0.01: **; 0.0001<*p*-value<0.001: ***; *p*-value<0.0001: ****). L-BH, 5-benzyl-L-hydantoin; BVH, 5-bromovinylhydantoin; L-IMH, 5-indolylmethyl-L-hydantoin, L-NMH 5-(2-naphthylmethyl)-L-hydantoin.

The transport protein Mhp1 from *Microbacterium liquefaciens* (Uniprot D6R8X8) transports 5-substituted hydantoins in a sodium-coupled mechanism (Suzuki and Henderson, 2006; Shimamura et al., 2010) (**Fig 2c**), and is a member of the nucleobase-cation-symport family of homologous proteins that also transport purines, pyrimidines, vitamins and related metabolites. The transporter has the LeuT-fold (Shimamura et al., 2010; Yamashita et al., 2005) and belongs to the amino acid-polyamine-organocation superfamily (Vastermark and Saier, 2014), which currently contains 20 families and 147,819 sequence entries, the substrates of which have mostly not been identified (Pfam CL0062). Mhp1 contains three cysteine residues, of which only one is readily accessible to reaction with *N*-ethylmaleimide (Calabrese et al., 2017). Purified Mhp1 in dodecyl-maltoside had an apparent T_m_ of 51.3 °C ± 0.6 °C (**Supplementary Fig. 4**) and to test the binding specificity, the ΔT_m_ was determined upon addition of different nucleobases and other compounds (**Fig. 2d**). The only stabilizing compounds were 5-indolylmethyl-L-hydantoin (ΔT_m_; 14.1 ± 1.2 °C), 5-benzyl-L-hydantoin (ΔT_m_; 13.6 ± 1.2 °C) and 5-bromovinylhydantoin (ΔT_m_; 11.9 ± 1.1 °C). A chemically related inhibitor of Mh1p, 5- (2-naphthylmethyl)-L-hydantoin (L-NMH) also led to a thermostability shift (ΔT_m_; 15.6 ± 1.7 °C). These results match the known substrate and inhibitor specificity of Mhp1 (Simmons et al., 2014), showing that the assay could identify substrates from a set of closely related compounds without false positives.

To validate the method further, we also screened the mitochondrial ADP/ATP carrier from *T. thermophila* (UniProt G2QNH0) (King et al., 2016) (**Fig. 2e**). This transport protein belongs to the mitochondrial carrier family (MCF), which is involved in the translocation of chemically diverse compounds across the mitochondrial inner membrane, using uniport, symport or antiport modes of transport (Kunji, 2012). Currently, there are 89,340 different sequence entries from 831 different species in the database (Pfam PF00153). In humans, the MCF is the largest solute carrier family with 53 different members (SLC25), but the substrate specificity of only half of them has been defined (Kunji, 2012). We screened the thermostability of purified AAC using a library of mitochondrial compounds (**Supplementary Fig. 5**). In the presence of ATP, ADP, and dADP the population was stabilized, showing ΔT_m_ values of 7.2 ± 0.2, 6.0 ± 0.5 and 1.8 ± 0.2 °C, respectively (**Fig. 1b** and **2f**). Some other compounds, mostly structurally related nucleotides, also stabilized the protein, but with significantly smaller shifts (**Fig. 2f**). For each type of nucleotide, the di- and tri-phosphate species showed larger shifts than the monophosphate forms, similar to the preference of ATP and ADP over AMP (**Fig. 1d** and **2f**), showing that the assay can also identify functional groups that are important contributors to substrate binding.

### Substrate screening of a mitochondrial carrier from *Tetrahymena thermophila*

To see whether this method can be used to identify candidate substrates for a previously uncharacterized transporter, we performed a high-throughput screen on a purified mitochondrial carrier from the thermophilic ciliate *Tetrahymena thermophila* (UniProt I7M3J0). The carrier is phylogenetically related to the yeast mitochondrial carrier that transports inorganic phosphate in symport with a proton into the mitochondrial matrix for ATP synthesis (Runswick et al., 1987), but its substrates have not been identified experimentally. The population of carriers had an apparent T_m_ of 56.0 ± 0.8 °C and was screened against a library of 132 different mitochondrial compounds at pH 6.0 (**Fig. 3**). The highest shift in the T_m_ of the population was observed for phosphate (ΔT_m_; 4.6 ± 0.4 °C), followed by glyoxylate (ΔT_m_; 3.6 ± 0.8 °C), acetyl-phosphate (ΔT_m_; 3.2 ± 0.5 °C) and phosphoenolpyruvate (ΔT_m_; 2.3 ± 0.2 °C) (**Supplementary Fig. 6**), whereas small shifts were observed for nucleotides.

**Figure 3.**
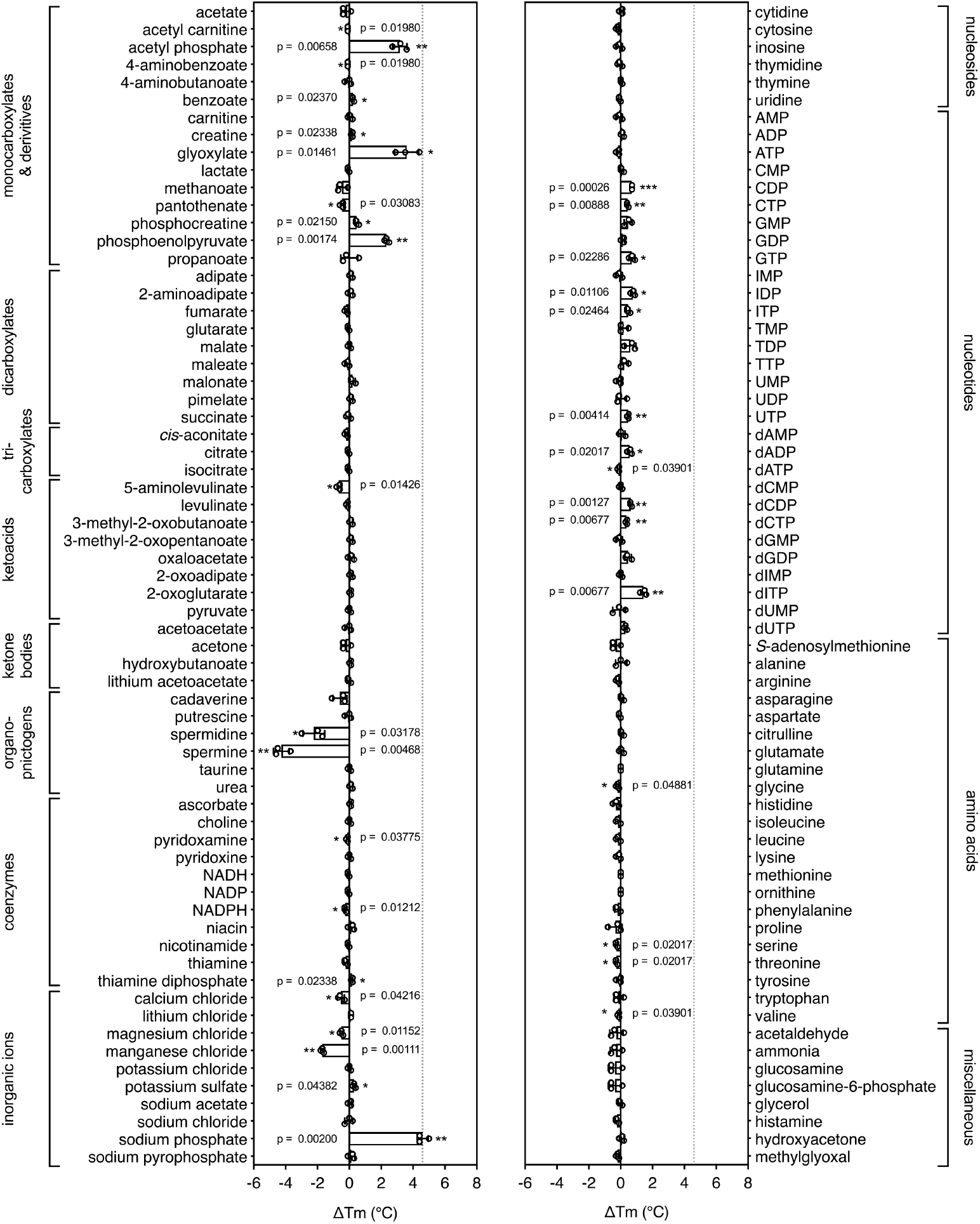
Identification of substrates for a mitochondrial phosphate carrier from the thermophilic ciliate *Tetrahymena thermophila*. Purified carrier (1 µg) in lauryl maltose neopentyl glycol was incubated at pH 6.0 with each library compound separately and the ΔT_m_ was determined, which is the difference between the apparent melting temperatures in the presence and absence of the tested compound. The data are represented by the average and standard deviation of three independent assays. The significance tests were performed as described in the legend to **Fig. 2.**

These compounds either contain phosphate as a functional group or resemble the structure of phosphate, such as glyoxylate (**Supplementary Fig. 7**). Thus, this assay can be used to narrow down substantially the number of potential substrate candidates from a library of unlabeled compounds.

### The effect of coupling ions

Secondary active transporters are widespread and often use the electrochemical gradient of protons or other ions to drive the uptake of substrates against their concentration gradient. Coupling ions are also recognized by transporters through specific interactions, often directly associated with the binding of substrates (Kunji and Robinson, 2010; Krishnamurthy et al., 2009).

5-Benzyl-L-hydantoin is transported by Mhp1 in symport with sodium ions (Suzuki and Henderson, 2006; Shimamura et al., 2010) and thus we tested whether the temperature shifts of Mhp1 induced by substrate binding were ion-dependent. When different ions were tested, only sodium ions induced a large substrate stabilization, whereas relatively small stabilizations were observed in the presence of potassium, calcium and magnesium ions (**Fig. 4a**).

**Figure 4.**
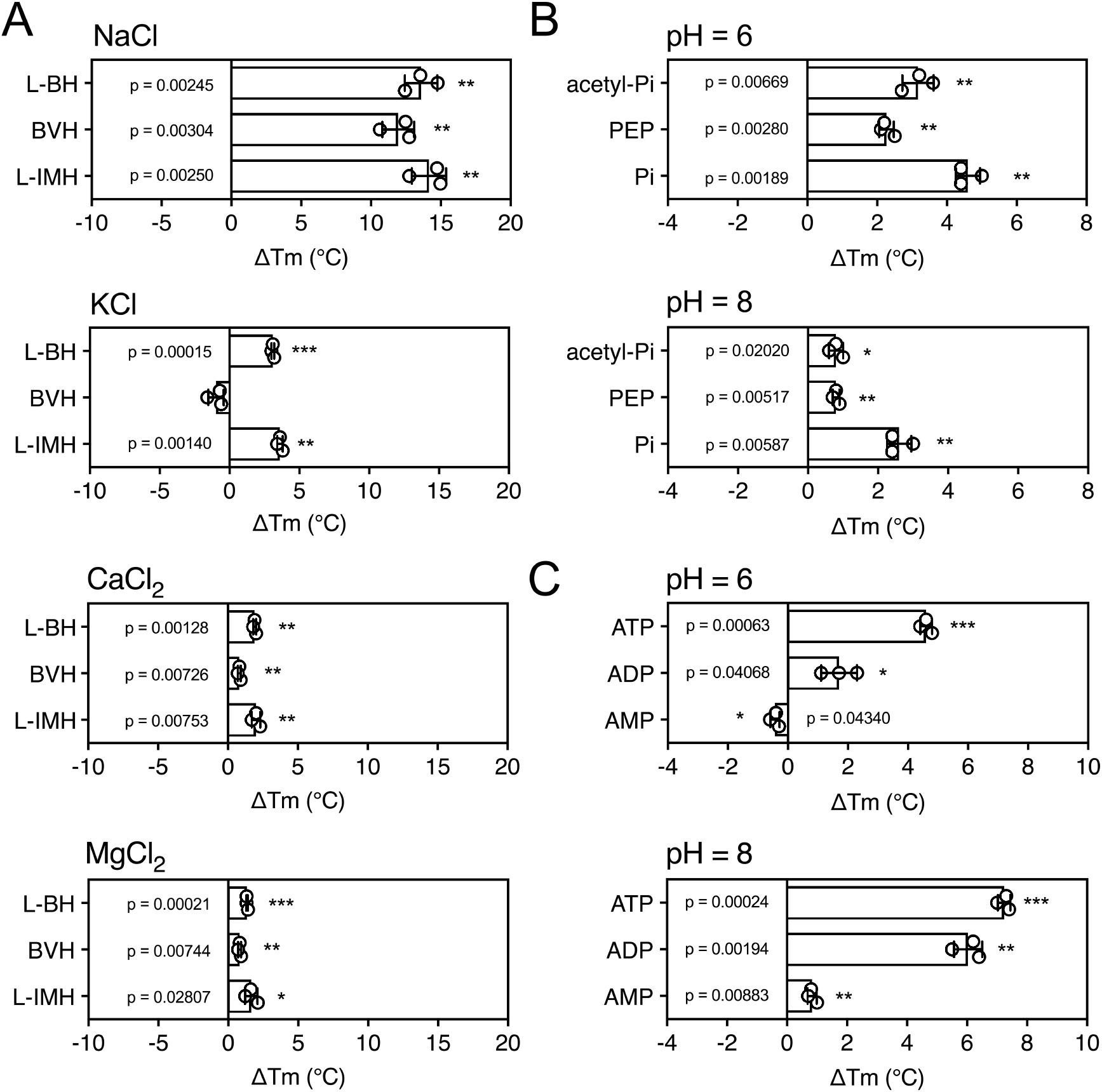
The effect of coupling ions on the stabilization of transporters by substrate binding. **(a**) Thermostability shifts of Mhp1 (3 µg) induced by binding of hydantoins in the presence of NaCl, KCl, CaCl_2_ and MgCl_2_. L-BH: 5-benzyl-L-hydantoin, BVH: 5-bromovinylhydantoin, L-IMH: 5-indolmethyl-L-hydantoin. (**b**) Thermostability shifts of the phosphate carrier (1 µg) induced by binding of phosphate-containing compounds at pH 6.0 and 8.0. Acetyl-Pi: acetyl-phosphate, Pi: phosphate. (**c**) Thermostability shifts of the (2 µg) induced by ADP and ATP binding at pH 6.0 and 8.0, using AMP as control.

Mitochondrial phosphate carriers are proton-driven, coupling phosphate translocation to proton symport (Kunji and Robinson, 2010). To investigate whether proton-dependency of substrate binding could be inferred from the assay, the shift in thermal stability by substrate binding was determined at pH 6.0 and 8.0. The pH can affect the reaction of maleimide groups of CPM with free thiols and the stability of the protein itself, but the ΔT_m_ value shows the effect of pH on substrate binding only. In both cases a substrate-induced stabilization was observed, but the effect was larger in pH 6.0 than in pH 8.0 (ΔT_m_ 4.6 ± 0.4 °C and 2.6 ± 0.4 °C, respectively) (**Fig. 4b**).

As a control, we carried out the same experiment with the mitochondrial ADP/ATP carrier, which is not a proton symporter (**Fig. 4c**). The opposite effect was observed; substrate binding at pH 8.0 led to a bigger thermostability shift than at pH 6.0. The predicted substrate binding site of AAC has three positively charged residues that interact with the negatively charged phosphate groups of ADP and ATP (Kunji and Robinson, 2006; Robinson and Kunji, 2006). ATP has pKa values of 0.9, 1.5, 2.3 and 7.7, whereas ADP has pKa values 0.9, 2.8 and 6.8. This means that the substrates are more negatively charged at pH 8.0 compared to pH 6.0, which will enhance their binding to the positively charged residues of the substrate binding site, leading to an increase in thermostability. In the case of the phosphate carrier the situation is very different. Phosphate has pKa values of 2.2, 7.2 and 12.3, and the predicted binding site contains two positively charged residues to neutralize the two negative charges on phosphate. However, in addition, the binding site contains a negatively charged glutamate, which has a pKa of ˜4.0. Only when this residue is protonated can binding of the negatively charged phosphate to the site occur (Kunji and Robinson, 2010), which occurs more readily at pH 6.0 than at pH 8.0, explaining the difference in thermostability. These results show that substrate binding stabilizes the transporters, but that this effect is enhanced further by the presence of coupling ions, providing testable hypotheses on energy coupling mechanisms.

## Discussion

We have shown that thermostability assays can be used to study the interaction of substrates with transporters. This is not intuitive, as substrates only bind transiently and rather weakly in the transport cycle. Still, for diverse types of transport proteins, which differ in structure and transport mechanism, a shift in thermostability was observed in the presence of specific substrates. The shift is observed at concentrations well above the apparent Km of transport, as under these conditions a larger part of the protein population is rescued from unfolding by binding of the substrate, increasing the overall number of interactions.

These studies highlight the different contributions these assays can make to studying the properties of transport proteins. First, they can be used to limit the number of potential substrate candidates from libraries of unlabeled compounds, providing important clues about the substrate specificity (**Fig. 2** and **3**). In the tested cases, the assay correctly identified the substrates from a set of chemically related compounds, including stereo-isoforms. Still, these candidates need to be tested by transport or other types of assays, as the compounds could potentially be inhibitors or regulators. However, no prior knowledge is required and the number of radioactive compounds that need testing is highly reduced, meaning that it is easier and cheaper to establish a robust transport assay. Second, the assays can be used to extend the substrate specificity of known transport proteins, as was shown for GalP with D-glucosamine (**Fig. 2b**). Third, they can be used to identify functional groups that are key to substrate binding, as shown for AAC, where nucleotide di- and tri- phosphates bound more tightly than nucleotide monophosphates (**Fig. 2f**). In the case of mitochondrial phosphate carriers, the common properties of compounds, such as the phosphate group, may lead directly to the most probable candidates. Fourth, this assay can provide important clues on the involvement of coupling ions in substrate binding, as shown for cases where transport is driven by sodium ion or proton symport (**Fig. 4**).

This method has major advantages over the traditional methods of identification. It is highly reproducible and can be performed in a high-throughput manner, allowing screening of about 100 compounds per machine per hour. The assay requires a relatively small amount of protein per assay (micrograms), depending on the number of buried cysteine residues. Another advantage is that the unfolding curves are themselves important quality checks, as they indicate that the protein is folded and therefore competent to bind substrate.

There are also some limitations, as only proteins that have buried thiols can be used, although accessible cysteines can be tolerated as they only raise the base line of the assay without interfering with the unfolding curve. Thiol-containing compounds, such as cysteine or glutathione, cannot be screened as the probe reacts with them directly. However, changes in the fluorescence of environmentally-sensitive dyes or endogenous tryptophans can be used as alternatives to CPM. Furthermore, the assay requires purified and stable transporter in detergent, although partially purified protein might be sufficient. Advances in bacterial (King et al., 2015; Ward et al., 1999), yeast (Routledge et al., 2016), insect (Contreras-Gomez et al., 2014), and mammalian (Goehring et al., 2014; Andrell and Tate, 2013) expression systems have allowed many transporters to be expressed in folded and functional states. Also, robust purification methods and novel classes of ‘stabilizing’ detergents, such as the neopentyl glycol maltoside detergents (Chae et al., 2010), have allowed even poorly expressed transporters to be purified in the small quantities required for these studies.

Even though the main purpose of this study was to identify potential substrates of transporters, the assays may also be used in drug discovery or in the identification of substrates, inhibitors and regulators of any other soluble or membrane proteins with occluded cysteines.

## Materials and Methods

### Materials

Chemicals were obtained from Sigma Aldrich and Thermo Fisher Scientific (USA). Nickel NTA and sepharose beads were purchased from Qiagen (USA). All enzymes were provided by New England BioLabs (USA). Lipids were purchased from Avanti Polar Lipids (USA) and detergents from Anatrace (USA). The hydantoin compounds were a kind gift of Marta Sans, Maria Kokkinidou and Arwen Pearson (University of Hamburg).

### Protein expression

For expression of mitochondrial ADP/ATP carrier of *Thermothelomyces thermophila* and the putative phosphate carrier of *Tetrahymena thermophila* in yeast mitochondria, gene constructs were designed to contain an N-terminal tag composed of eight histidine residues followed by a Thr-Ser-Glu-Asp linker and an Ile-Glu-Gly-Arg Factor Xa protease cleavage site. The genes were cloned into pYES3/CT vector (Invitrogen, USA) with a constitutively active promoter (*pMIR* promoter of the *S. cerevisiae* phosphate carrier). The plasmids were transformed into *S. cerevisiae* WB.12 (MATα ade2-1 trp1-1 ura3-1 can1-100 aac1::LEU2 aac2::HIS3) and W303-1B (MATα leu2-3,112 trp1-1 can1-100 ura3-1 ade2-1 his3-11,15) strains respectively. Cells were grown in a 50-litre fermenter after which mitochondria were prepared (Kunji and Harding, 2003).

For expression of GalP in *E. coli* the promoter region and the *galP* gene, which was modified to encode six histidine residues at the C-terminus of the protein, were cloned into plasmid pBR322 to form plasmid pGP1, which was transformed into the galactose/glucose transport-deficient host strain JM1100 (*ptsG ptsM ptsF mgl galP Hfr Δhis gnd thyA galK)* (Henderson et al., 1977). Cells were grown in basal salts medium supplemented with 30 mM glycerol, 20 µg/ml thymine, 80 µg/ml histidine and 15 µg/ml tetracycline in a fermenter (30- or 100-liter scale). The gene *hyuP* from *M. liquefaciens*, modified to encode six histidine residues at the C-terminus, was cloned into plasmid pTTQ18 (Stark, 1987; Suzuki and Henderson, 2006). The His_6_-tagged Mhp1 hydantoin transport protein was expressed in *E. coli* BL21 (DE3) grown in M9 medium supplemented with 20 mM glycerol, 20 mM NH_4_Cl, 100 μg/ml of carbenicillin, 0.2% w/v casamino acids, 2 mM MgSO_4_.7H_2_O, 0.4 mM CaCl_2_.2H_2_O, using induction by IPTG in 100-liter fermenter cultures. In all cases after harvesting the intact cells preparations were made of inner membranes (Ward et al., 1999), which were stored at –80^°^C in Tris-HCl buffer pH 7.5 until used for purification of each individual protein.

### Protein purification

The mitochondrial ADP/ATP carrier and phosphate carrier were purified using the detergents dodecyl-maltoside and lauryl maltose neopentyl glycol, respectively, using established procedures (King et al., 2016).

For the purification of His-tagged GalP, *E. coli* membranes were solubilized in buffer containing 20 mM Tris-HCl pH 8, 20 mM imidazole, 300 mM NaCl, 20% glycerol and 1% dodecyl-maltoside for 1 h at 4 °C with gentle mixing to a final protein concentration of 2.5 mg/ml. After centrifugation (108,000 × g, 1 h, 4 °C), the supernatant was mixed with pre-washed nickel-NTA resin and purified by immobilized nickel affinity chromatography (1 ml resin per 37 mg of total protein; Superflow, QIAGEN) for batch-binding affinity chromatography for 1 h at 4 °C. Non-specific proteins were removed with buffer containing 20 mM Tris-HCl pH 8, 20 mM imidazole, 150 mM NaCl, 10% glycerol and 0.02% dodecyl-maltoside, after which His-tagged GalP was eluted with buffer containing 20 mM Tris-HCl pH 8, 200 mM imidazole, 100 mM NaCl, 5% glycerol and 0.02% dodecyl-maltoside. Imidazole was then removed by passing the sample through pre-equilibrated PD-10 desalting column (GE Healthcare). Protein samples were snap-frozen and stored in liquid nitrogen.

The purification of Mhp1 followed the same procedure as the GalP purification with the following modifications. Membranes were solubilized for 2 h and incubated with Nickel NTA resin for 2 h. The wash buffer contained 10 mM Tris-HCl pH 8, 20 mM imidazole, 10% glycerol and 0.05% dodecyl-maltoside and the elution buffer contained 10 mM Tris-HCl pH 8, 200 mM imidazole, 2.5% glycerol and 0.05% dodecyl-maltoside.

### Transport assays

Robotic transport assays were performed using a Hamilton MicroLab Star robot (Hamilton Robotics Ltd, UK). For GalP transport assays, *E. coli* cells (GalP-expressing strain pGP1/JM1100 and JM1100 control strains) were diluted to OD_600_ of 20 in MES buffer (5 mM MES pH 6.6, 150 mM KCl) and incubated at room temperature with 20 mM glycerol for 10 minutes to be energized prior to being loaded into the wells of a MultiScreenHTS+ Hi Flow-FB (pore size = 1.0 μm, Millipore, USA). Transport assays were initiated by addition of 100 μl of buffer containing 5 μM [^14^C]-galactose (American Radiolabeled Chemicals, 0.20 GBq/mmol) or 5 μM [^14^C]-glucosamine (American Radiolabeled Chemicals, 0.204 GBq/mmol). For the competition assay, radiolabeled substrate was mixed with a 4,000-fold excess of the competitor compound prior to the assay. The transport was stopped after 0, 10, 20, 30, 45, 60, 150 s and, 5, 7.5 and 10 min incubation times with 200 μl ice-cold buffer and the samples were filtered and washed as above. Levels of radioactivity were measured by adding 200 μl MicroScint-20 (Perkin Elmer) and measured with a TopCount scintillation counter (Perkin Elmer).

### Preparation of the mitochondrial compound library

To identify mitochondrial metabolites every enzyme in the KEGG metabolic database (Kanehisa et al., 2017) was evaluated for mitochondrial localization using the MitoMiner database of mitochondrial localization evidence (Smith and Robinson, 2016). A wide range of data were considered including large-scale experimental evidence from GFP tagging and mass-spectrometry of mitochondrial fractions, mitochondrial targeting sequence predictions, immunofluorescent staining from the Human Protein Atlas (Thul et al., 2017), and annotation from the Gene Ontology Consortium (The Gene Ontology, 2017). Ensembl Compara allowed these data to be shared across human, mouse, rat and yeast homologs (Zerbino et al., 2018). Once a list of enzymes with probable mitochondrial localization was collated, KEGG was used to produce a corresponding list of potential mitochondrial compounds. Additional candidates were taken from a computer model of mitochondrial metabolism that manually partitioned metabolites on either side of the mitochondrial inner membrane (Smith and Robinson, 2011). Compounds were dissolved in PIPES buffer (10 mM PIPES pH 7.0, 50 mM NaCl) to a final concentration of 25 mM or in dimethyl sulfoxide to a final concentration of 100 mM. pH was adjusted if necessary and the stocks were frozen at −80 °C.

### Analysis of the total number of identified transport protein substrates in different species

The TransportDB database contains a large number of organisms for which the transporter complement has been identified via a bioinformatic pipeline, along with substrate predictions for the transporters characterized (Elbourne et al., 2017). To acquire the figures in Supplementary Table 1, a Perl script driving an SQL query of the underlying MySQL database to TransportDB was developed to quantify those transporters where no prediction could be made for the substrate for the listed species. The number of transporters with unassigned specificity represents a minimal number, as the search procedure uses sequence similarity to characterised transporters as a criterion. However, the substrate specificity needs to be determined experimentally, as a single mutation in the binding site can profoundly change substrate recognition.

### Thermostability shift assay and library screening

To determine stability, purified protein (typically 1-3 μg) was mixed with 20 μg/ml of the thiol-reactive fluorophore 7-diethylamino-3- (4’-maleimidylphenyl)-4-methylcoumarin (CPM) and the fluorescent intensity was measured over a 25-90 °C range of temperature using a rotatory qPCR multi sample instrument (Rotor-Gene Q, Qiagen, the Netherlands). Following an initial 18 °C pre-incubation step of 90 s, the temperature was ramped 1 °C every 15 s, with a 4-s wait between readings, which is equal to a ramp rate of 5.6 °C/min. The excitation and emission wavelengths were 460 nm and 510 nm, respectively. Five mg/ml stocks of CPM prepared in dimethyl sulfoxide were diluted to 0.1 mg/ml and equilibrated in assay buffer for one hour at room temperature in the dark before addition to the protein sample. The assay buffer was usually the buffer in which the protein was purified or in a similar buffer with a different pH (MES for pH 6.0, HEPES for pH 8.0) or a different concentration of other ions. Data analyses and apparent melting temperatures (T_m_, the inflection point of a melting temperature) were determined using software supplied with the instrument.

For GalP experiments, 500 mM sugar stocks were made in MilliQ water (Merck-Millipore, USA) as 10 times stocks. For Mhp1 experiments, 100 mM compound stocks were made in dimethyl sulfoxide as 50 times stocks.

### Data analyses and representation

Statistical analyses were performed using Microsoft Excel with the inbuilt function of two-tailed, two-sample unequal variance Student’s *t*-test. The average apparent T_m_ of ‘no ligand’ control samples (three technical repeats within each Rotor-Gene Q run) was subtracted from the apparent T_m_ measured for each compound addition in the same run. This assay was performed with three or five biological repeats using independent batches of purified protein. The null hypothesis of the *t*-test was that the observed ΔT_m_ for each compound was not significantly different from zero.

## Acknowledgments

This work was supported by grant MC_UU_00015/1 of the Medical Research Council, UK. H.M. gratefully acknowledges the Cambridge Commonwealth, European and International Trust for support of her PhD studies. PJFH thanks the Leverhulme Trust for an Emeritus Research Fellowship (Grant number EM-2014 –045). The fermenters and allied equipment for protein production in Leeds were funded by the BBSRC (MPSI BBS/B/14418), the Wellcome Trust (JIF 062164/Z/00/Z) and the University of Leeds. The hydantoin derivatives were a kind gift from Marta Sans, Maria Kokkinidou and Arwen Pearson (University of Hamburg).

## Author contributions

H.M. and M.S.K. contributed protein purification and thermostability shift assays; H.M., M.S.K. contributed transport assays; S.M.P. and D.S. contributed cell culturing; A.C.S. contributed the mitochondrial compound library; L.D.H.E and I.T.P contributed bioinformatics analyses of substrate characterization; H.M., M.S.K., P.J.F.H. and E.R.S.K. contributed the design of the experiments, analyses of the data and writing of the manuscript.

## Supplementary Table

**Supplementary Table 1.** Status of predictions of transport protein substrate specificity in representative archaeal, eubacterial and eukaryotic species (April 2018).

| Genus | Species | Transporter<br>Proteins | Proteins<br>with no<br>predicted<br>substrate | Percentage |
| --- | --- | --- | --- | --- |
| *Acinetobacter* | *baumannii* | 391 | 75 | 19% |
| *Arabidopsis* | *thaliana* | 1279 | 311 | 24% |
| *Caenorhabditis* | *elegans* | 669 | 179 | 27% |
| *Campylobacter* | - | 197 | 37 | 19% |
| *Drosophila* | *melanogaster* | 662 | 105 | 16% |
| *Enterococcus* | *faecium* | 392 | 44 | 11% |
| *Escherichia* | *coli* | 638 | 120 | 19% |
| *Halobacterium* | - | 208 | 10 | 5% |
| *Helicobacter* | *pylori* | 140 | 33 | 24% |
| *Homo* | *sapiens* | 1472 | 102 | 7% |
| *Klebsiella* | *pneumoniae* | 879 | 164 | 19% |
| *Neisseria* | *gonorrhoeae* | 172 | 32 | 19% |
| *Oryza* | *sativa* | 1285 | 165 | 13% |
| *Pseudomonas* | *aeruginosa* | 703 | 147 | 21% |
| *Saccharomyces* | *cerevisiae* | 343 | 126 | 37% |
| *Salmonella* | - | 605 | 113 | 19% |
| *Shigella* | - | 534 | 95 | 18% |
| *Staphylococcus* | *aureus* | 348 | 41 | 12% |
| *Streptococcus* | *pneumoniae* | 307 | 50 | 16% |
| *Sulfolobus* | - | 205 | 18 | 9% |
| *Vibrio* | *cholerae* | 458 | 82 | 18% |
| Total |  | 11886 | 2048 | 18% |

## Supplementary Figures

**Supplementary Figure 1.**
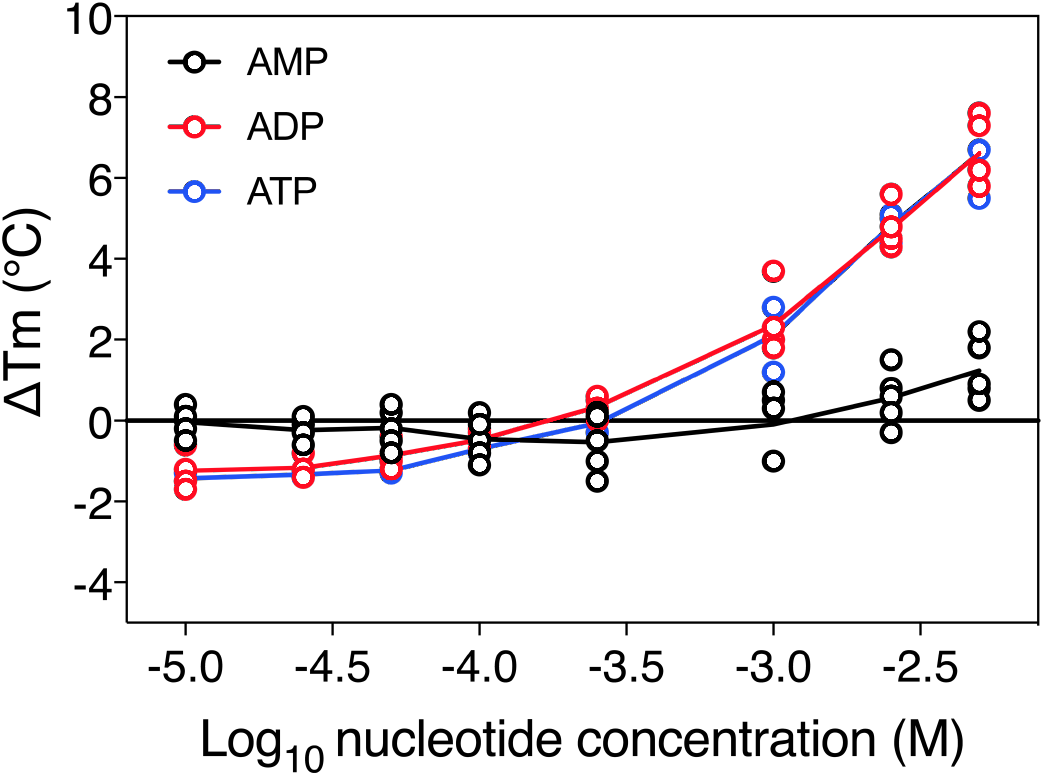
The substrate concentration-dependency of the apparent melting temperature of purified mitochondrial ADP/ATP carrier. The apparent melting temperature (T_m_) was determined from the peak in the derivative of the unfolding curve for different concentrations of AMP, ADP, and ATP in 20 mM HEPES pH 8.0, 100 mM NaCl, 0.1 % dodecyl-maltoside, 0.1 mg ml^-1^ tetraoleoyl cardiolipin. The data are represented by the average and standard deviation of three biological repeats.

**Supplementary Figure 2.**
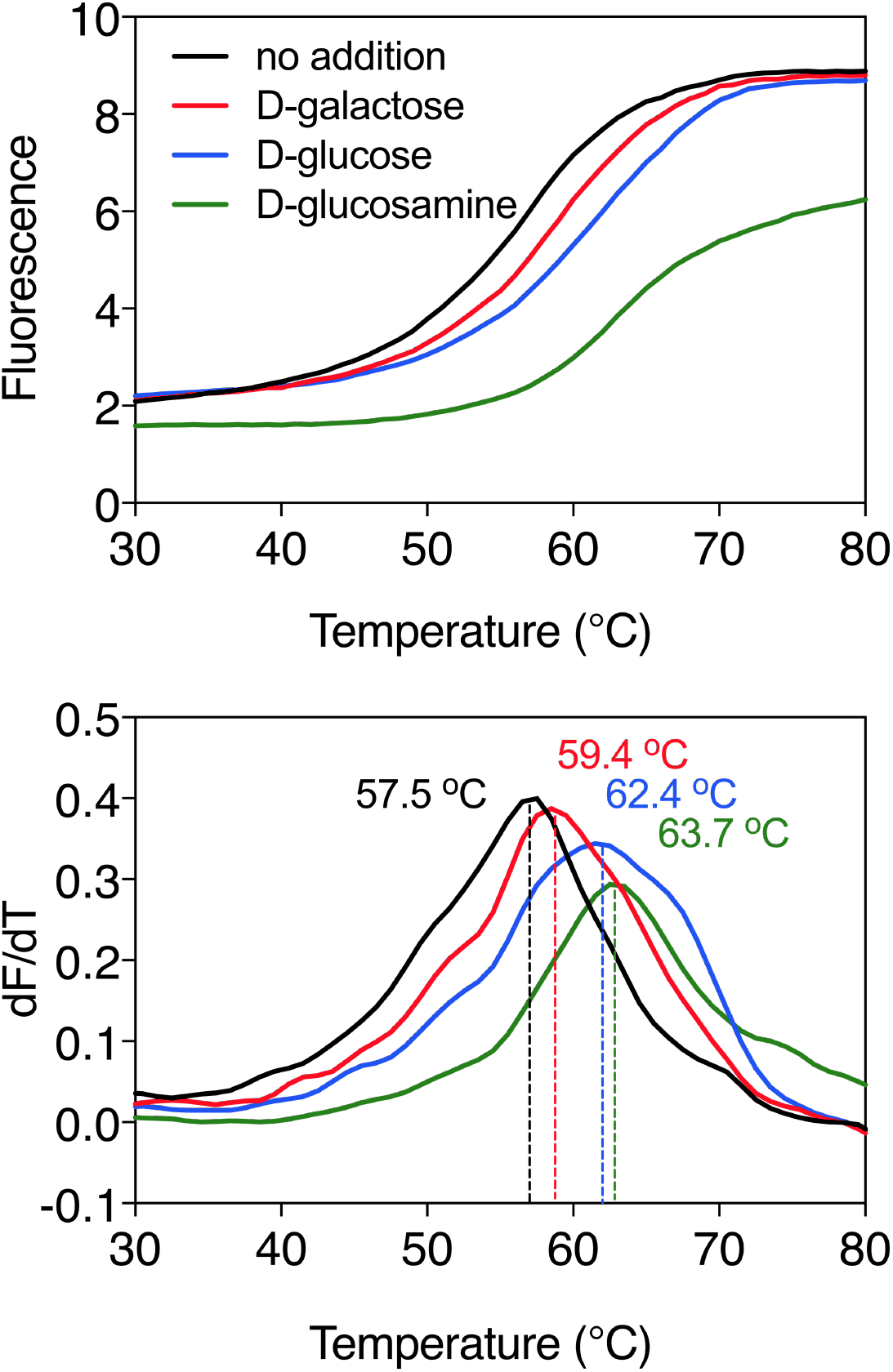
Substrate-induced stabilization of the galactose transporter GalP. Typical thermostability profiles of purified GalP (**top**) and their derivatives (dF/dT; **bottom**) in the absence and presence of stabilizing substrates D-galactose (red trace), D-glucose (blue trace) and glucosamine (green trace), which was also causing a significant shift. The apparent melting temperature (T_m_) is the peak in the derivative of the unfolding curve (dF/dT) and is indicated for each curve. For each reaction, 2.5 μg purified protein was assayed in the presence of 50 mM compound. The apparent melting temperature reported in the text is from five independent protein purifications.

**Supplementary Figure 3.**
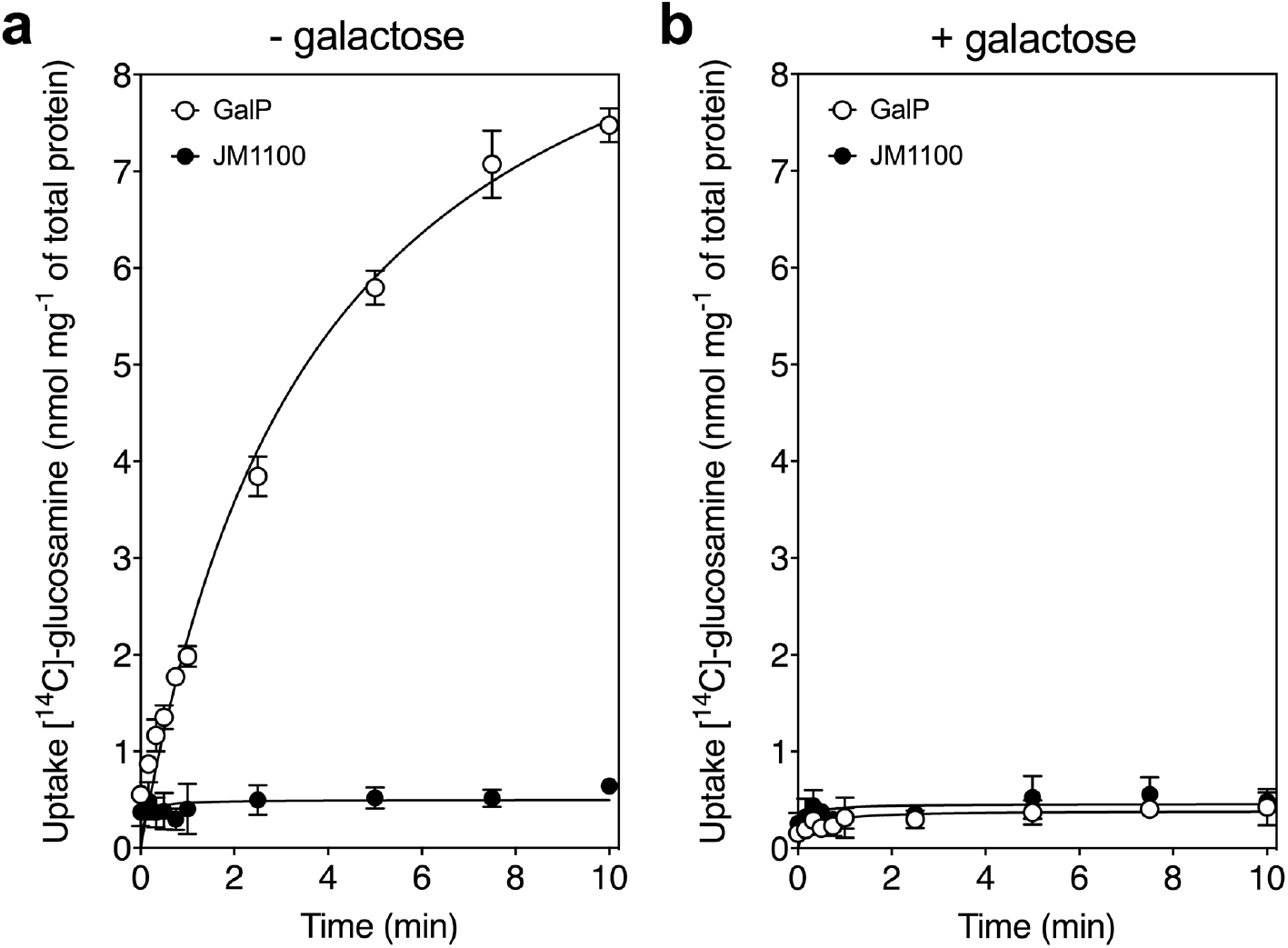
Glucosamine is a transported substrate of the galactose transporter GalP. Transport assays with [^14^C]-labeled D-glucosamine in whole cells of the GalP-expressing plasmid pGP1 in the host *E. coli* strain JM1100 (open circles) and control JM1100 without plasmid (closed circles). Transport in (**a**) the absence or (**b**) presence of 4,000-fold excess of unlabeled D-galactose as described in the “Transport assays” section. The data are represented by the average and the standard deviation of four technical repeats.

**Supplementary Figure 4.**
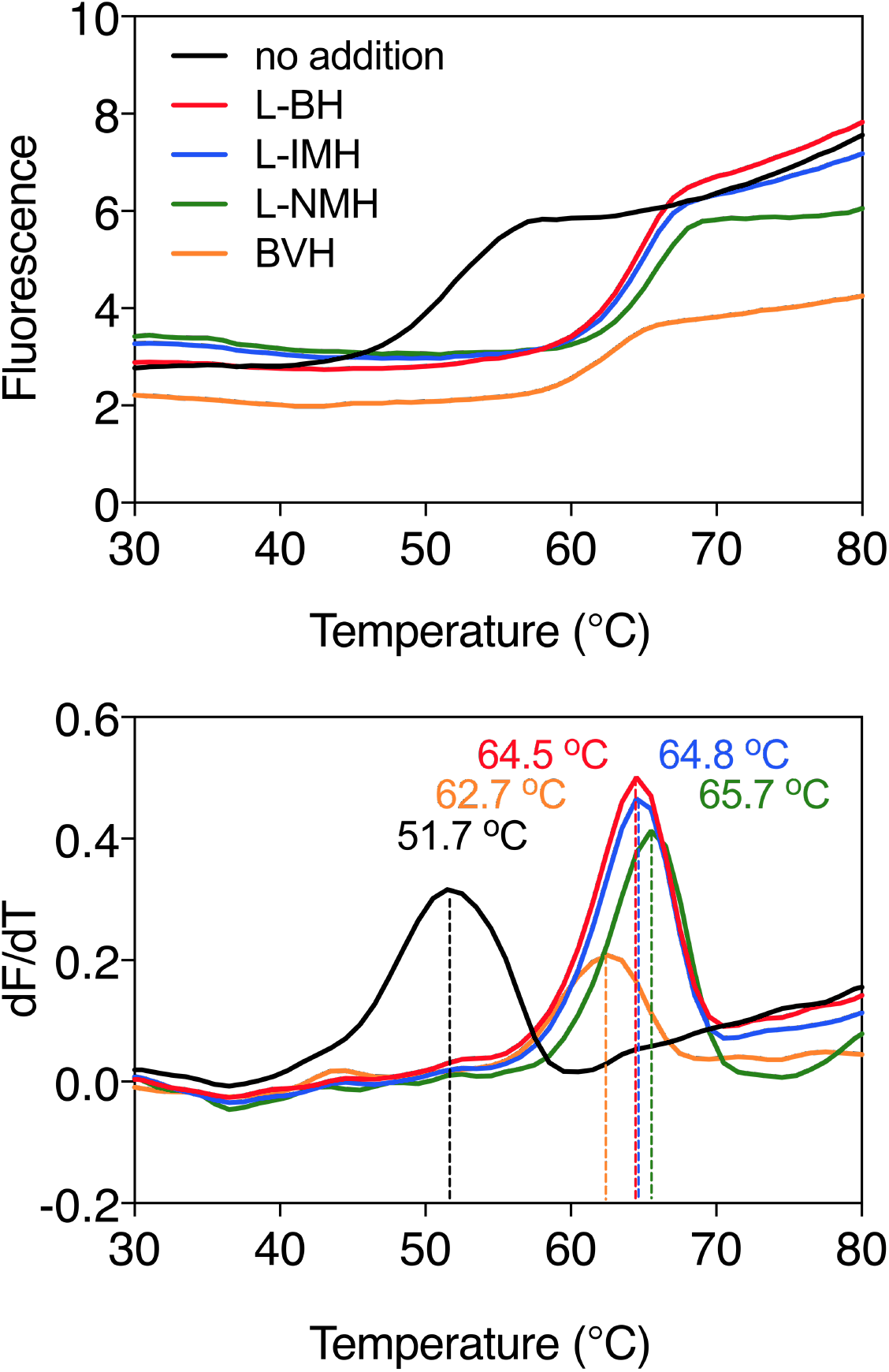
Ligand-induced stabilization of Mhp1 in the presence of sodium ions. Typical thermostability profiles of purified Mhp1 (**top**) and their derivatives (dF/dT; **bottom**) in the absence or presence of stabilizing compounds 5-benzyl-L-hydantoin (L-BH, red trace), 5-indolylmethyl-L-hydantoin (L-IMH, blue trace), 5-(2-naphthylmethyl)-L-hydantoin (L-NMH, green trace) and 5-bromovinylhydantoin (BVH, orange trace). The apparent melting temperature (T_m_) is the peak in the derivative of the unfolding curve (dF/dT) and is indicated for each curve. For each reaction, 2.5 μg purified protein was assayed in the presence of 140 mM NaCl and 2 mM compound in buffer containing 10 mM Tris-HCl pH 8.0, 2.5% glycerol and 0.05% dodecyl-maltoside. The apparent melting temperature reported in the text is from three independent protein purifications.

**Supplementary Figure 5.**
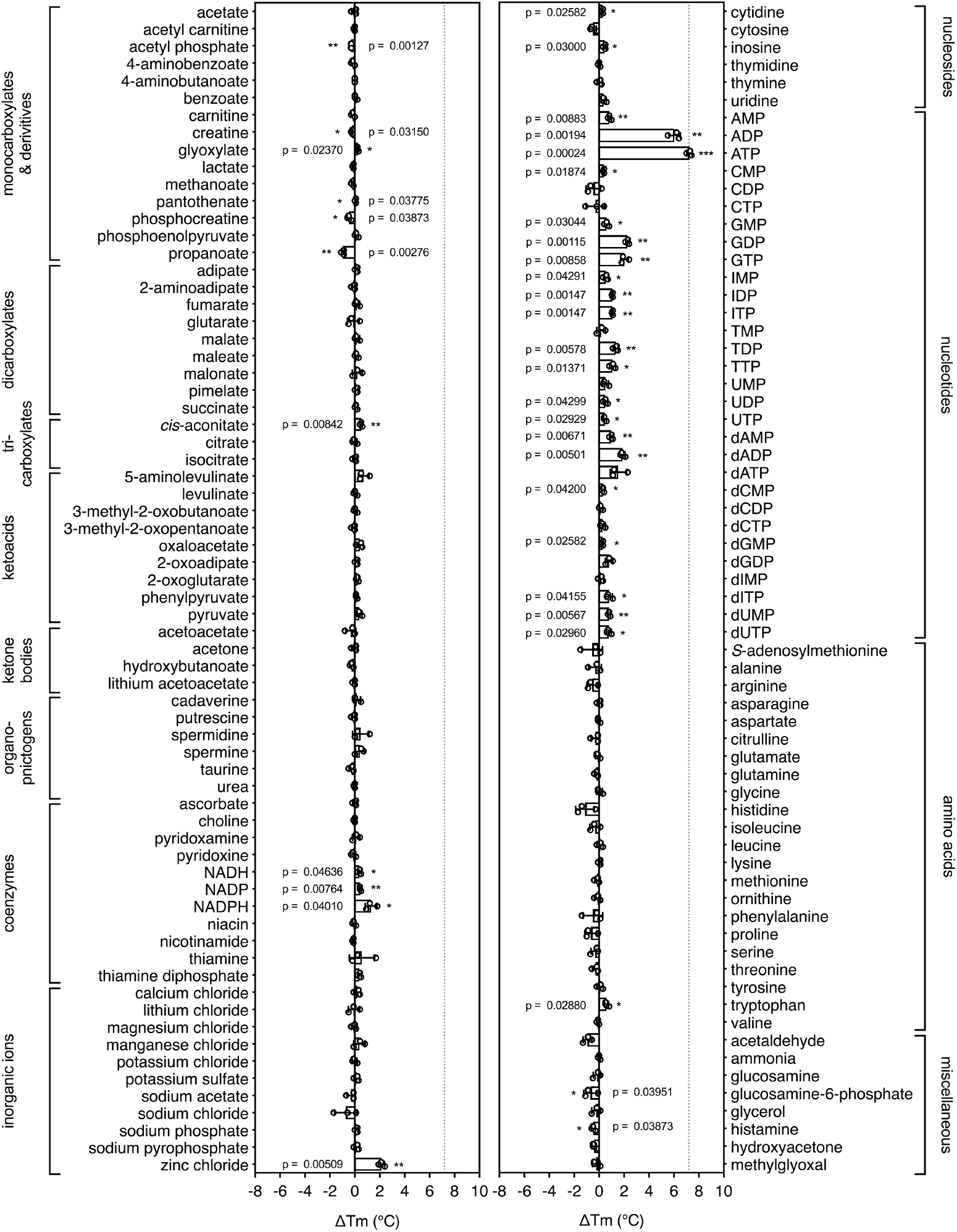
Substrate screening of the mitochondrial ADP/ATP carrier. Purified ADP/ATP carrier in dodecyl-maltoside was separately mixed with 2.5 mM of library compounds in 20 mM HEPES pH 8.0, 100 mM NaCl, 0.1 % dodecyl-maltoside, 0.1 mg ml^-1^ tetraoleoyl cardiolipin and subjected to a melting regime. The shifts in the apparent T_m_ (see Online Methods) were recorded and are shown as bars. The data are represented by the average and standard deviation of three independent assays. The significance tests were carried out as in **Figure 2**.

**Supplementary Figure 6.**
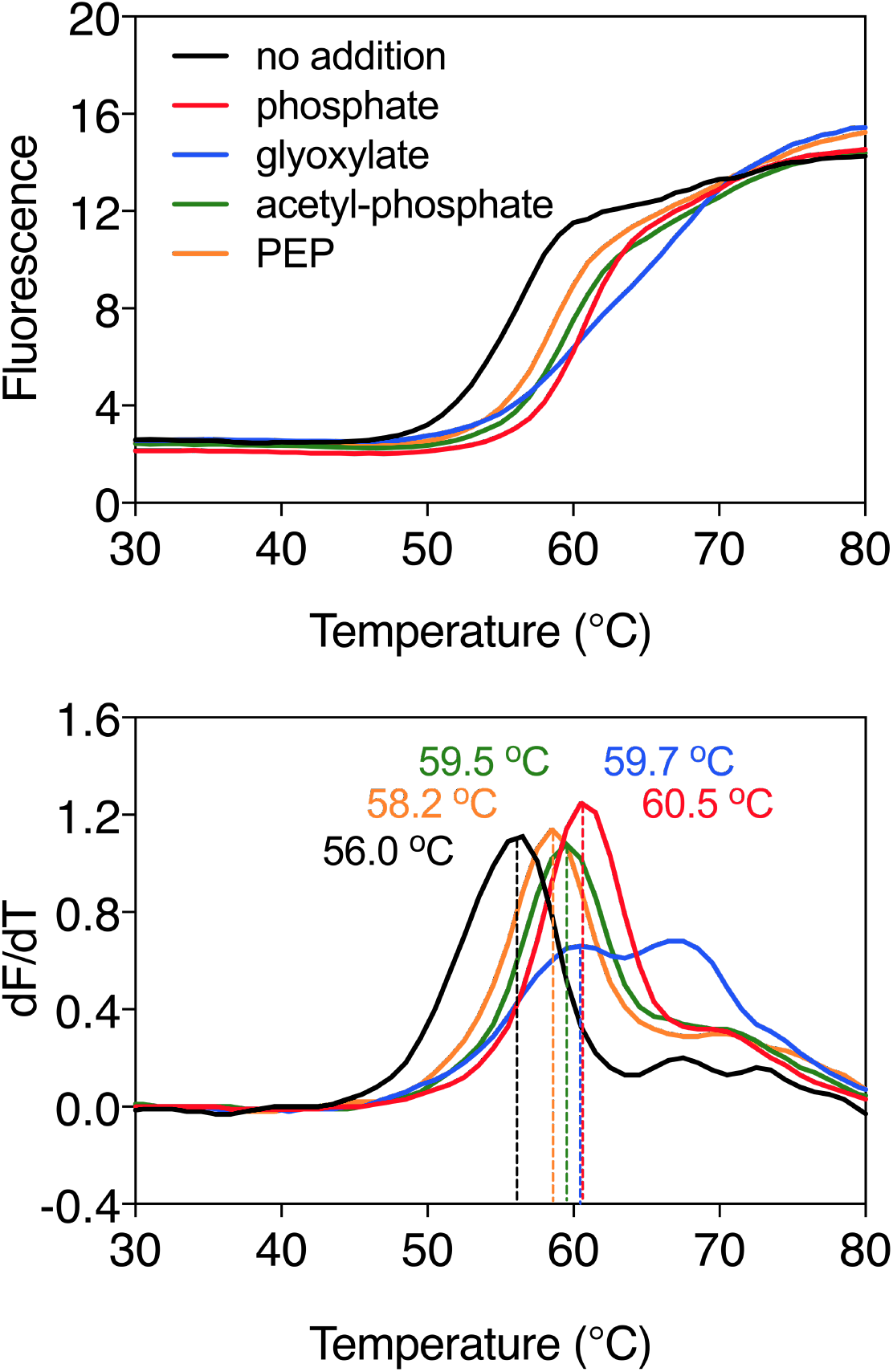
Substrate-induced stabilization of the mitochondrial phosphate carrier. Typical thermostability curves of purified phosphate carrier (**top**) and their derivatives (dF/dT; **bottom**) in the absence and presence of stabilizing compounds phosphate (PEP, red trace), glyoxylate (blue trace), acetyl-phosphate (green trace) and phosphoenolpyruvate (orange trace). The apparent melting temperature (T_m_) is the peak in the derivative of the unfolding curve (dF/dT) and is indicated for each curve. For each reaction, 2 μg purified protein was assayed in the presence of 2.5 mM compound in 20 mM MES pH 6.0, 100 mM NaCl, 0.1 % lauryl maltose neopentyl glycol, 0.1 mg ml^-1^ tetraoleoyl cardiolipin. The apparent melting temperatures reported in the text are from three independent protein purifications.

**Supplementary Figure 7.**
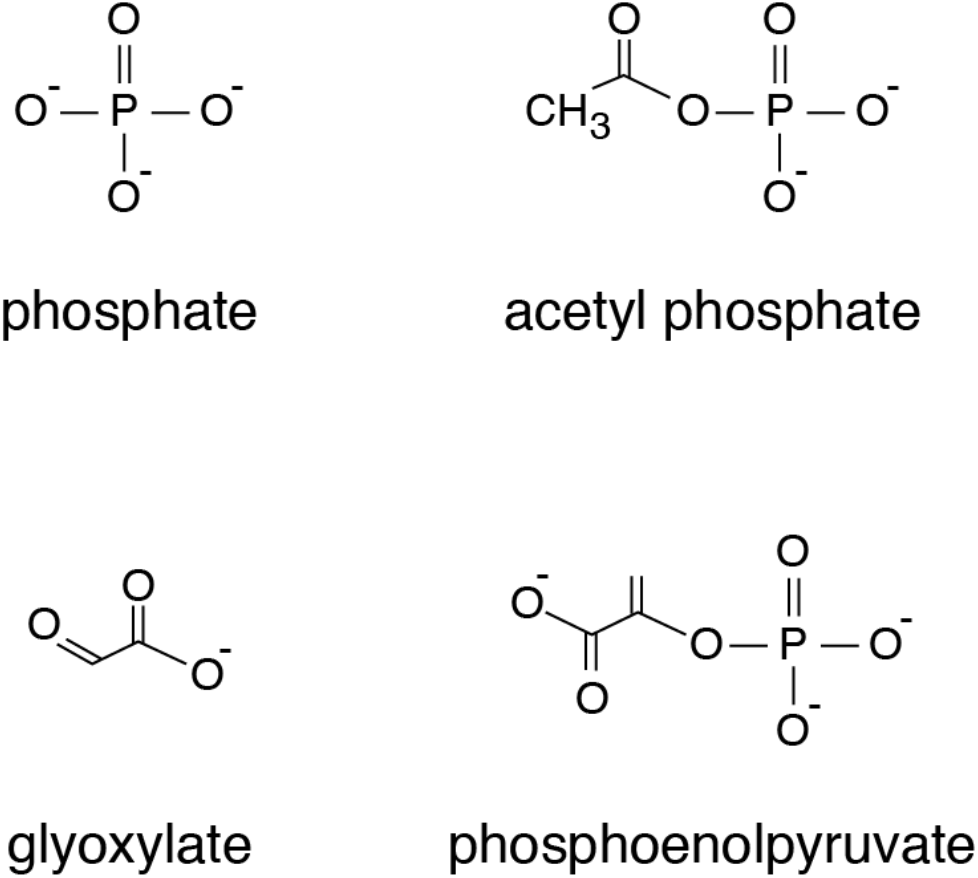
Chemical structures of compounds that showed significant thermostability shifts with the mitochondrial phosphate carrier.

